# Hierarchical refinements of cis-regulatory inputs improve scalable gene expression prediction

**DOI:** 10.64898/2026.05.31.729151

**Authors:** Qinhu Zhang, Meng Xing, Qi Liao, Zhipeng Li, De-Shuang Huang

## Abstract

Deciphering the relationships between cis-regulatory elements (CREs) and target gene expression has long been a challenging problem in molecular biology. However, predicting gene expression from hundreds of candidate cis-regulatory elements (cCREs) requires models that scale to long, noisy inputs while retaining interpretable regulatory structure. Existing Transformer-based approaches typically attend over all nucleotides and all surrounding cCREs, diluting causal signals when hundreds of elements compete for limited model capacity. Here we introduce a **<u>t</u>**wo-**<u>s</u>**tage **<u>s</u>**elective **<u>f</u>**ramework (TSSF) that performs hierarchical refinements: nucleotide-level masking within each cCRE, followed by cCRE-level selection around each gene, implemented with information-bottleneck priors and a fully Transformer-based architecture. Across 70 human cell types and tissues, TSSF and lightweight variants improve expression prediction and enhancer–gene prioritization relative to strong baselines, including on cross-cell-line and cell-type-specific benchmarks. Prediction-stratified analysis motivates a distance-decay prior that aligns attention with long-range regulatory geometry, and chromatin-contact augmentation improves recovery of distal links. Motif analyses of high-confidence predictions recover proximal and distal regulatory programs, supporting mechanistic interpretability. TSSF offers a general strategy for scalable, interpretable modeling of high-dimensional regulatory inputs in genomics.

## 1. Introduction

Predicting gene expression from genomic and epigenomic information is central to understanding how cellular identity and phenotype are determined^1^. Gene expression is primarily controlled by *cis*-regulatory elements (CREs)—non-coding sequences such as enhancers and promoters that regulate when and where genes are transcribed. With millions of candidate cis-regulatory elements (cCREs) present in mammalian genomes, linking these elements to their target genes is crucial for interpreting genetic variation, identifying disease mechanisms, and connecting genome-wide association signals to disease-causing genes^2^. Consequently, accurate computational prediction of gene expression from cCREs would provide a powerful, scalable complement to costly and time-consuming experimental assays.

Conventionally, gene expression has been measured directly using hybridization-based or sequencing-based technologies such as microarrays and RNA-seq. At the same time, regulatory relationships have been inferred through expression quantitative trait locus (eQTL) mapping^3,4,^ chromatin conformation capture^5,6,^ and targeted assays like STARR-seq and CRISPR-based perturbations^7^. Although these approaches have yielded valuable insights, they suffer from several limitations: experimental protocols are costly and low-throughput; RNA-seq analysis is sensitive to preprocessing and pipeline choices, leading to non-negligible variability in expression estimates across studies; and many methods are restricted to specific tissues or cell types, making it difficult to generalize regulatory logic at scale. Thus, there remains a need for computational frameworks that can predict gene expression from sequence and epigenomic assays without relying on exhaustive experimental characterization of each cCRE–gene pair.

Deep learning has substantially advanced this problem. Convolutional and hybrid models such as DeepSEA^8^, Expecto^9^, and Basenji2^10^ learn local motif syntax and long-range sequence context, while Enformer^11^ extends receptive fields to ∼100 kb using Transformer. More recent cCRE-centric frameworks, including GraphReg^12^, CREaTor^13^, EPInformer^14^, ScPGE^15^, and Seq2Exp^16^—integrate epigenomic tracks and, in some cases, 3D contact maps to relate discrete regulatory elements to transcriptional output. Despite these advances, a shared computational bottleneck remains: for each gene, models must process hundreds of cCREs, each represented by hundreds of nucleotides and multiple epigenomic channels. Standard Transformer encoders process all nucleotides within cCREs and all candidate regulatory elements uniformly, without explicitly distinguishing causal from redundant or weakly informative inputs—attention is learned over very large, heterogeneous, noisy input spaces, which can dilute key regulatory signals when hundreds of cCREs are considered per gene.

From a modeling perspective, gene regulation is naturally hierarchical: only a subset of nucleotides within an element carry functional motif syntax, and only a subset of surrounding cCREs actively influence expression in a given cell type. Explicitly enforcing this hierarchy—rather than expecting a single attention layer to discover it implicitly—should improve both predictive accuracy and post-hoc prioritization of functional enhancer–gene pairs. Information bottleneck (IB) principles provide a natural formalism: learn compressed masks that retain maximally informative inputs for gene expression prediction task while suppressing redundant variation^17,18.^

Here we propose a **<u>t</u>**wo-**<u>s</u>**tage **<u>s</u>**elective **<u>f</u>**ramework (TSSF) that operationalizes hierarchical refinements in a fully Transformer-based architecture. In stage 1, TSSF uses nucleotide-level representations to generate masks for selecting the most highly correlated nucleotides within each cCRE. In stage 2, it uses cCRE-level representations to generate masks for selecting the subset of cCREs most strongly correlated with gene expression. These masks are parameterized as Beta distributions with IB-inspired KL regularization, enabling differentiable training via reparameterization. By performing hierarchical refinements at the nucleotide and cCRE levels, TSSF effectively reduces input redundancy and concentrates model capacity on regulatory features most predictive of transcription. Furthermore, we introduce two variants of TSSF, namely TSSF-signal and TSSF-nobeta, respectively. Both variants achieve performance comparable to TSSF at a lower cost in terms of model complexity. We benchmark TSSF on 70 human cell types and tissues against CREaTor, EPInformer, ScPGE, and Seq2Exp, and evaluate (i) cross-cell-line and cross-species transfer, (ii) cell-type-specific expression, (iii) cCRE–gene prioritization on curated benchmarks, and (iv) CRISPR-validated enhancers at the KLF1 and GATA1 loci. Beyond predictive performance, we show that prediction-stratified analysis motivates a distance-decay prior, that chromatin contacts preferentially improve distal element recovery, and that high-confidence cCREs recover known TF programs. Together, these results position hierarchical refinements as a general computational strategy for high-dimensional regulatory modeling, with immediate applications in scalable functional genomics.

## 2. Results

### 2.1 Hierarchical refinement architecture

TSSF refines regulatory representations in two coupled stages before expression prediction. Stage 1 learns nucleotide-level masks within each discretized cCRE from sequence and DNase-seq (TSSF-signal uses DNase alone); stage 2 selects cCREs from multimodal epigenomic profiles and feeds refined representations to a cCRE Transformer that outputs gene expression levels. TSSF-nobeta removes explicit Beta masking and relies on standard attention to identify important nucleotides and cCREs. Specifically, as shown in Figure 1, (i) we extract discrete cCREs surrounding genes from the whole genome and arrange them in order of their positions; (ii) we convert DNA sequences into index vectors and extract epigenomic signals from corresponding epigenomic tracks, forming sequence features and chromatin features, respectively; (iii) TSSF generates a Beta distribution from DNA sequences and a Beta distribution from DNase-seq signals, respectively. By combining these two Beta distributions, we obtain a joint Beta distribution 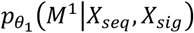 to identify key nucleotides; (iv) TSSF generates another Beta distribution 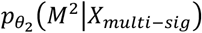 from multimodal epigenomic profiles to identify key cCREs; (v) all representations obtained through the two-stage refinement are used by a cCRE Transformer to predict gene expression levels (GPL).

**Figure 1:**
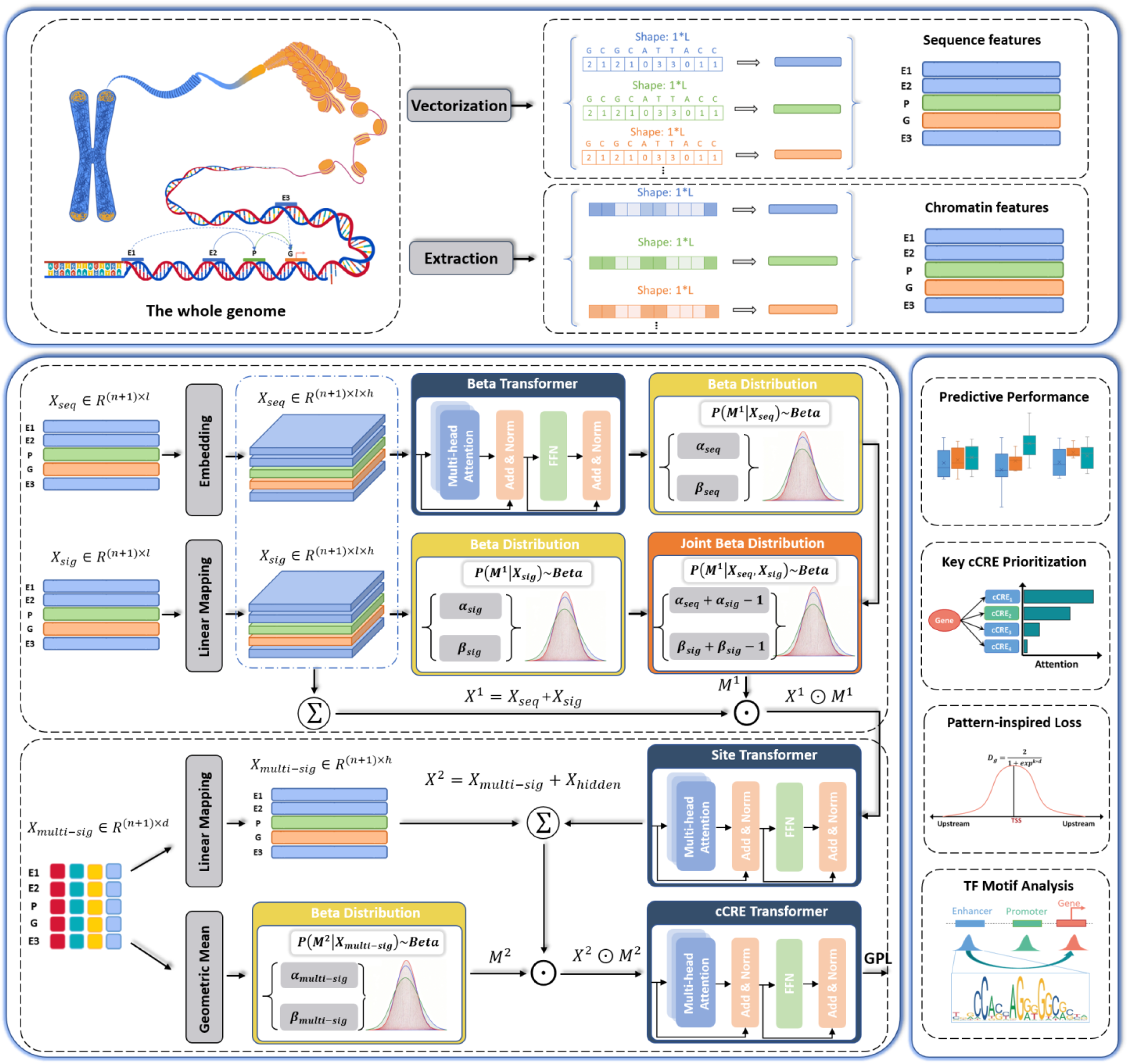
Schematic overview of TSSF. TSSF is a fully Transformer-based framework that performs hierarchical refinements in two stages: first, it uses nucleotide-level representations to generate masks for selecting the most highly correlated nucleotides within each cCRE; second, it uses cCRE-level representations to generate masks for selecting the subset of cCREs most strongly correlated with gene expression. During data construction, DNA sequences are vectorized into a two-dimensional matrix *X*_*seq*_; DNase-seq signals are correspondingly extracted from the epigenomic tracks to form a two-dimensional matrix *X*_*sig*_; in addition, various types of epigenomic profiles are converted into a two-dimensional matrix *X*_*multi*−*sig*_. During model computation, TSSF uses the nucleotide-level representations (*X*_*seq*_ and *X*_*sig*_) to generate a joint Beta distribution 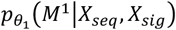 to screen for key nucleotides, and utilizes the cCRE-level representations (*X*_*multi*−*sig*_) to generate a Beta distribution 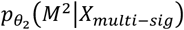 to screen for key cCREs; finally, the representations obtained through the two-stage refinement are employed to predict gene expression levels (GPL). During model evaluation, beyond predictive performance, the ability to prioritize functional enhancer–gene interactions, the prior-inspired loss enhancement, and the recovery of TF motifs are systematically explored.

Through a comprehensive comparison with various state-of-the-art methods, including predictive performance, the ability to prioritize active cCREs, cross-species prediction, and cell-type-specific gene prediction, we demonstrate the effectiveness of our proposed methods in gene expression prediction.

### 2.2 Systematic gains in expression prediction across 70 human profiles

We first asked whether hierarchical refinements improve genome-wide expression prediction. To this end, we adopted four state-of-the-art models | CREaTor^13^, EPInformer^14^, ScPGE^15^, and Seq2Exp^16^ | and compared them with our proposed two-stage frameworks (TSSF, TSSF-signal, and TSSF-nobeta). We quantified the performance of these models across multiple cell types and tissues using Pearson correlation coefficients (Pearsonr) and mean absolute error (MAE) metrics.

To comprehensively assess the predictive performance of these models, we used 70 human datasets covering a wide range of cell types and tissues. As shown in Figures 2a-b, our proposed two-stage frameworks outperform other competing methods, achieving the highest Pearsonr and the lowest MAE. Moreover, among the two-stage frameworks, both TSSF-signal and TSSF-nobeta achieve comparable performance to TSSF while incurring lower model complexity and computational resource costs. These differences likely reflect distinct input representations: (i) CREaTor and ScPGE encode DNA sequences and epigenomic tracks without hierarchical refinement, so nucleotide- and element-level redundancy in the raw inputs can dilute predictive signal. They also do not incorporate per-cCRE summaries of epigenomic activity (e.g., the average epigenomic signal for each cCRE), which TSSF uses in stage 2 to represent the global cCRE context. (ii) EPInformer relies on DNA sequence and a restricted set of epigenomic signals and does not explicitly model nucleotide-resolution accessibility; without DNase-seq, sequence alone may provide limited nucleotide-level regulatory information. (iii) Although Seq2Exp performs strongly, it does not aggregate per-cCRE epigenomic summaries to refine element-level representations before prediction. By contrast, our frameworks refine cCRE representations in two stages—nucleotides within each element, then elements around each gene—improving gene expression prediction. TSSF and TSSF-signal use Beta-distributed masks for both stages, whereas TSSF-nobeta replaces explicit Beta masking with learned attention while retaining the two-stage design.

**Figure 2:**
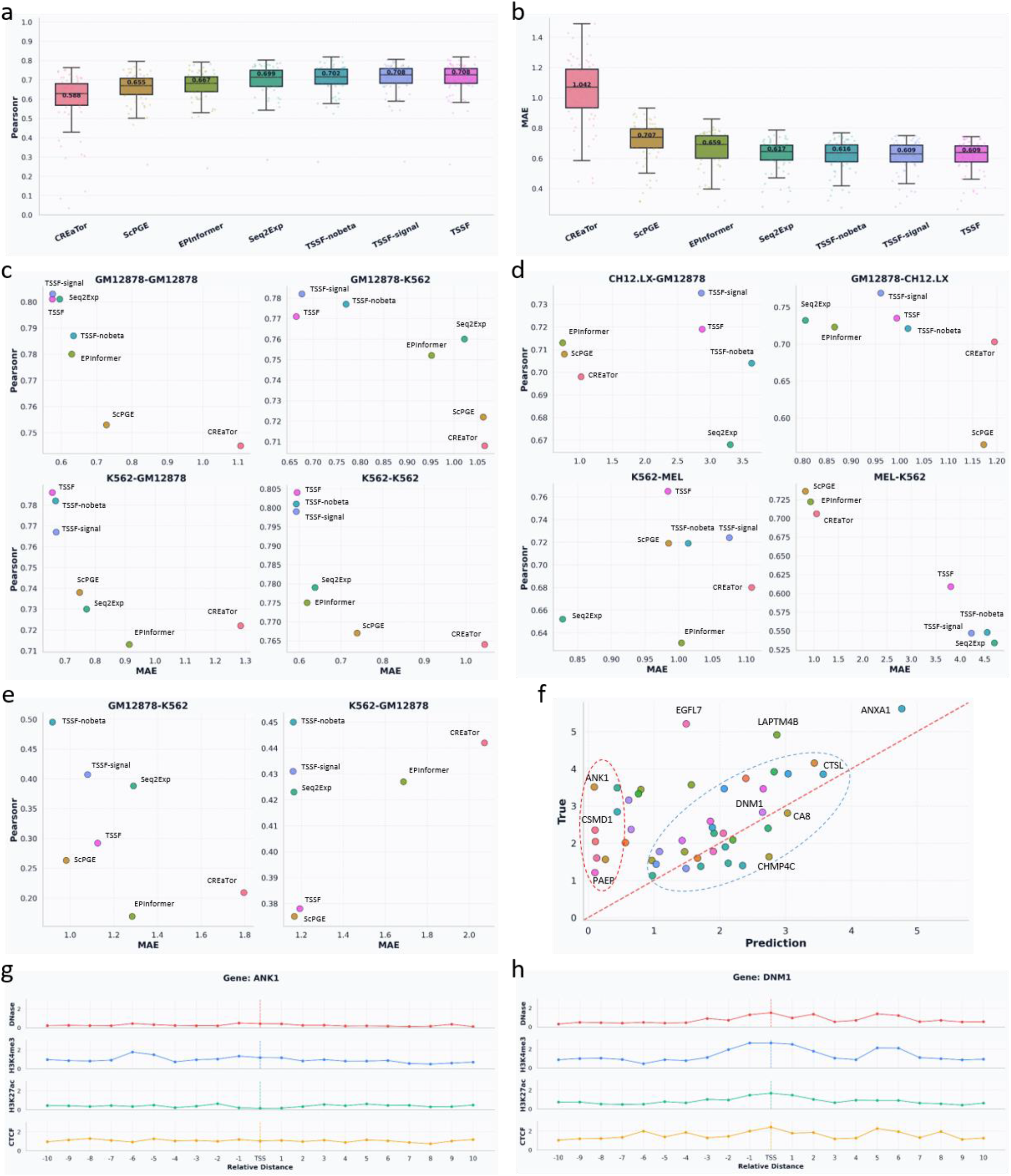
Overall performance of the two-stage frameworks. (a-b) Predictive performance (Pearsonr and MAE) of all methods across 70 human cell types and tissues. (c) Cross-cell-type predictive performance of all methods on GM12878 and K562 cell lines. (d) Cross-species predictive performance of all methods on human and mouse species. (e) Performance of all methods in predicting cell-type-specific gene expression. (f) Scatter plot of K562-specific gene using the Pearsonr metric. (g) Visualization of four epigenomic signals for an incorrectly predicted gene (*ANK1*). (h) Visualization of four epigenomic signals for a correctly predicted gene (*DNM1*).

Additionally, we conducted cross-cell-line and cross-species prediction experiments to further validate the performance of our proposed frameworks. In the cross-cell-line prediction experiments, we chose the GM12878 and K562 cell lines as the benchmark datasets. As shown in Figure 2c, our two-stage frameworks generally outperform other competing methods in predicting gene expression within the same cell line or across different cell lines. In the cross-species prediction experiments, we chose the mouse-derived CH12.LX and MEL cell lines as the corresponding benchmark datasets of the human-derived GM12878 and K562 cell lines. Specifically, we utilized the two-stage frameworks trained on the GM12878 and K562 cell lines to predict gene expression levels for the CH12.LX and MEL cell lines, respectively, and vice versa. As shown in Figure 2d, we observe an inconsistent result: our proposed frameworks generally outperform other methods in human-to-mouse cross-species prediction; however, they perform worse than other methods in mouse-to-human cross-species prediction. This observation suggests that these computational methods struggle to accurately predict gene expression levels across species due to significant heterogeneity among different species.

To further investigate the performance of our proposed frameworks in predicting cell-type-specific gene expression, we identified cell-type-specific genes from the K562 and GM12878 cells according to each gene’s log2FC. Subsequently, we used the trained models to predict cell-type-specific gene expression levels within the same cell line or across different cell lines. As shown in Figure 2e, our two-stage frameworks generally outperform other competing methods, and TSSF-nobeta achieves the best performance among the two-stage frameworks. Moreover, as shown in Figure 2f, we find that not all cell-type-specific genes can be accurately predicted; in particular, a small number of highly expressed genes are underestimated. To investigate this, we performed a visual analysis of four types of epigenomic signals on the cCREs surrounding two genes, one of which (*ANK1*) was incorrectly predicted and the other (*DNM1*) was correctly predicted. From the two examples (Figures 2g-h), we can see that DNase and H3K27ac signals near the incorrectly-predicted gene (*ANK1*) are very low, approaching 0, while values for other epigenomic signals are relatively high; on the contrary, the values for all types of epigenomic signals near the correctly-predicted gene (*DNM1*) are relatively high. These findings suggest that DNase and H3K27ac signals are key indicators for identifying active genes; if these data contain noise, this could lead to model misclassification.

### 2.3 Improved prioritization of functional cCRE–gene interactions

Linking candidate enhancers to their target genes through high-throughput experiments remains a significant challenge in terms of both time and cost. Against this backdrop, the use of computational methods to accurately prioritize candidate enhancer-gene interactions has become increasingly important. To evaluate the ability of the two-stage frameworks to capture true cCRE-gene interactions, we constructed three types of cCRE-gene interaction datasets, including an integrated enhancer-gene dataset, three GTEx datasets, and a Hi-ChIP dataset.

The integrated enhancer-gene dataset was collected from three publicly available sources, and divided into four groups based on distance: ‘<10kb’, ‘10-50kb’, ‘50-100kb’, and ‘>100kb’. We chose CREaTor, EPInformer, ScPGE, and ABC-pipeline^7^ as baseline models, and applied attention weights as well as augmented attention weights to prioritize key enhancers. As measured by the PRAUC metric (Figure 3a), our proposed frameworks perform better than other baseline methods in identifying valid cCRE-gene interactions, while TSSF performs best when using augmented attention weights (TSSF-ABC). Three eQTL fine-mapping datasets were collected from GTEx v10, involving ‘EBV-transformed Lymphocytes’, ‘Whole Blood’, and ‘Liver’, and similarly divided into four distance-based groups. As measured by the PRAUC metric (Figures 3b-d), our proposed frameworks generally outperform other baseline methods in identifying valid cCREs near genes, but perform poorly in identifying distal cCREs (‘>100kb’). For the Hi-ChIP interaction dataset, we similarly find that our two-stage frameworks perform better than other competing methods across different distance ranges (Figure 3e). The above results across three types of cCRE-gene interaction datasets demonstrate the effectiveness of our two-stage frameworks in prioritizing cell-type-specific cCREs.

**Figure 3:**
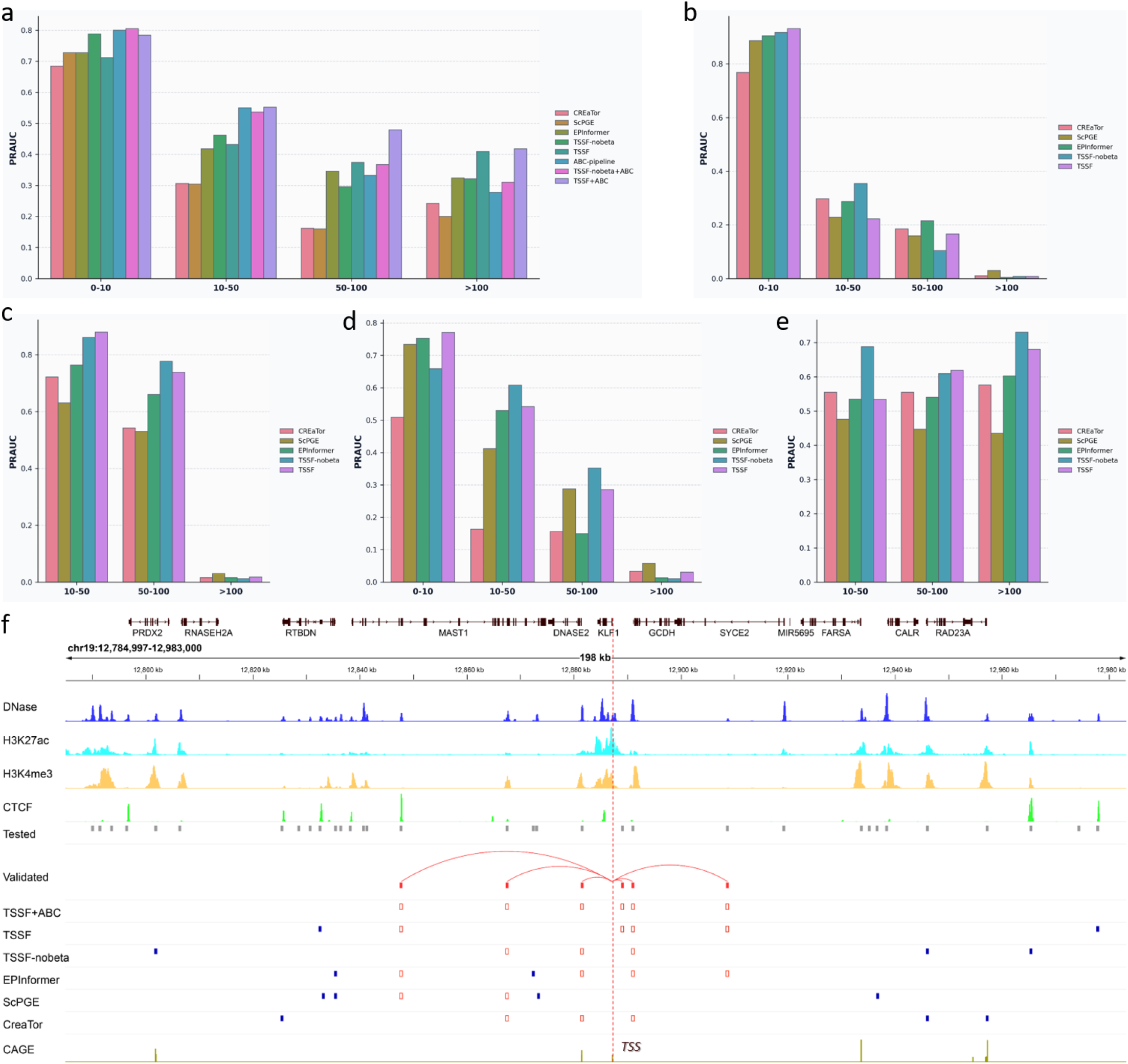
The ability of the two-stage frameworks to prioritize cell-type-specific cCREs. (a) Classification performance (PRAUC) of various methods across different distance groups in cCRE-gene interactions integrated from three publicly available sources. (b-d) Classification performance (PRAUC) of various methods across different distance groups in cCRE-gene interactions derived from three eQTL fine-mapping datasets: ‘EBV-transformed Lymphocytes’, ‘Whole Blood’, and ‘Liver’. (e) Classification performance (PRAUC) of various methods across different distance groups in cCRE-gene interactions derived from Hi-ChIP data. (f) Visualization of *KLF1* gene-active cCREs prioritized by multiple methods, where all tested cCREs are represented by solid grey boxes, validated cCREs by solid red boxes, and correctly-identified cCREs by red boxes.

To gain a deep understanding of all models’ ability to identify active enhancers, we focused on two important genes (*KLF1* and *GATA1*) in the K562 cell line. We selected all candidate enhancers within a 100 kb region surrounding these two genes as the test targets. These enhancers were evaluated using CRISPRi-FlowFISH assays, and only a few showed a significant effect on gene expression. To classify candidate enhancers as active or inactive, we prioritized all enhancers and selected top-*k* enhancers (where *k* is the number of validated enhancers) to calculate the true positive rate (TPR). For the *KLF1* gene (Figure 3f), there are six validated enhancers (solid red boxes) surrounding it, including three proximal enhancers and three distal enhancers. We observe that both TSSF and EPInformer identify four active enhancers (TPR=4/6), and outperform other methods in capturing active enhancers; in particular, TSSF+ABC (using augmented attention weights) achieves the highest TPR and discovers all active enhancers. For the *GATA1* gene (Supplementary Figure 1), there are five validated enhancers (solid red boxes) surrounding it, including two proximal enhancers and three distal enhancers. We find that TSSF identifies three active enhancers (TPR=3/5), and outperforms other methods in capturing active enhancers; similarly, TSSF+ABC (using augmented attention weights) achieves the highest TPR and discovers all active enhancers. These case studies demonstrate that hierarchical refinements sharpen attention on causally active elements beyond what global expression scores alone reveal.

### 2.4 Distance-decay prior aligns attention with regulatory geometry

Although various computational methods for predicting gene expression have been proposed, in-depth analyses of their predictions remain scarce. Such analyses are crucial for identifying unique patterns and providing feedback to optimize the models. To this end, we classified predictions from TSSF into true positives (TPs), false positives (FPs), true negatives (TNs), and false negatives (FNs) (see **Supplementary Note 2**). By visualizing multiple epigenomic signals of cCREs surrounding genes (Figure 4a and Supplementary Figures 2-3), we find that in TPs, these signals show an approximately normal-like distribution, reflecting that the regulatory effect of cCREs on target genes diminishes with increasing distance; whereas in TNs, the distribution is almost linear, with DNase and H3K27ac signals approaching zero. In particular, FPs exhibit a similar normal distribution, which explains why unexpressed genes are incorrectly predicted to be expressed.

**Figure 4:**
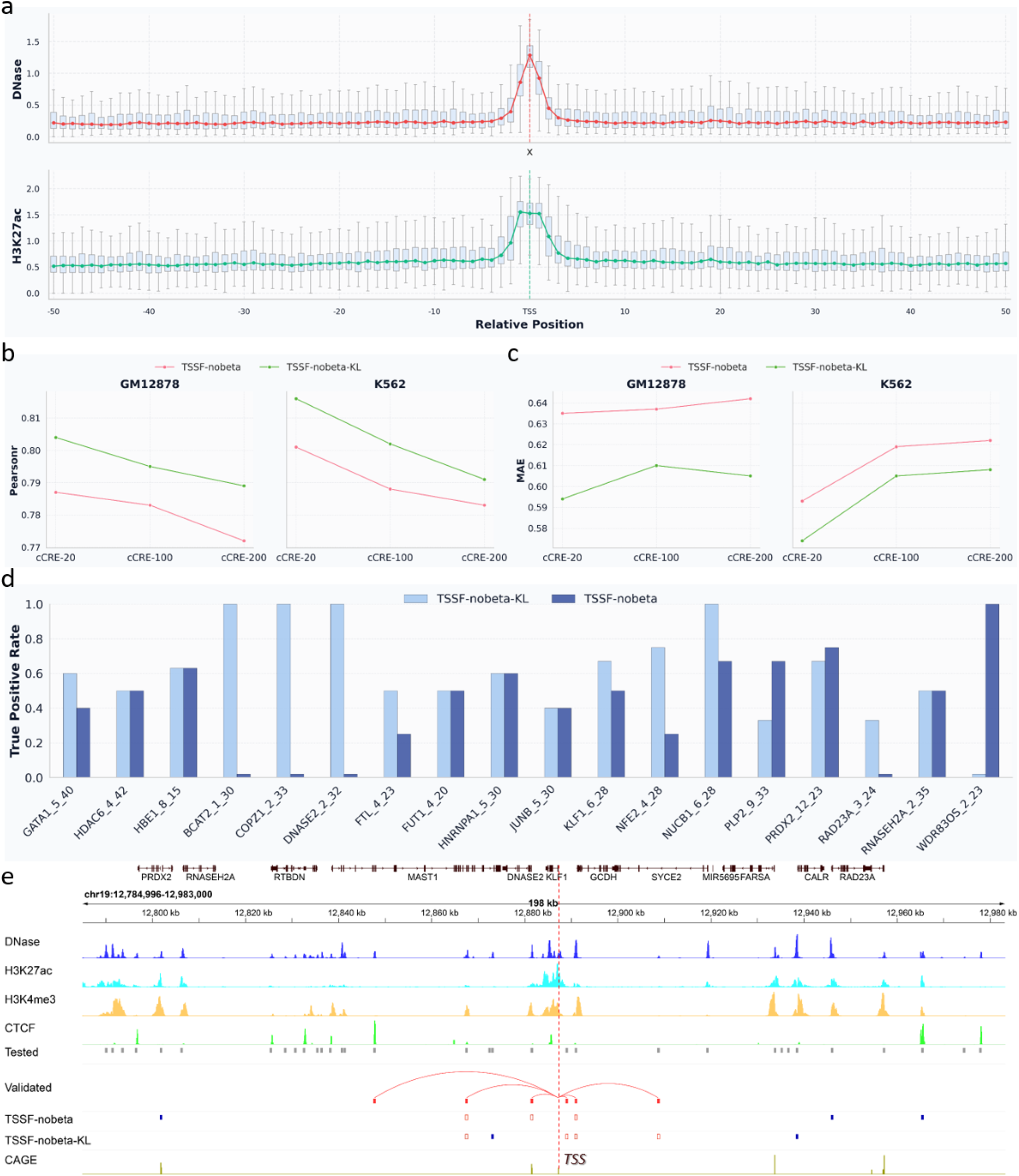
Prior-inspired loss improves the predictive performance. (a) A prior was found in TPs that shows an approximately normal-like distribution. (b-c) Performance comparison (Pearsonr and MAE) between TSSF-nobeta and TSSF-nobeta-KL. (d) True positive rates of TSSF-nobeta and TSSF-nobeta-KL in identifying genuine cCREs. (e) Visualization of *KLF1* gene-active cCREs prioritized by TSSF-nobeta and TSSF-nobeta-KL, where all tested cCREs are represented by solid grey boxes, validated cCREs by solid red boxes, and correctly-identified cCREs by red boxes.

Inspired by this prior, we designed a loss function based on the KL divergence between the prior and attention weights, where the prior is defined as an exponential decay function (see **Methods**). By aligning attention with regulatory geometry, we can guide the distribution of attention weights to converge toward the distribution of the decay function. As shown in Figures 4b-c, the predictive performance of TSSF-nobeta with KL is better than that of TSSF-nobeta, demonstrating the effectiveness of this prior-inspired loss. Furthermore, we applied models with this loss to prioritize cell-type-specific cCREs. Measured by the true positive rate (TPR), we observe that TSSF-nobeta with KL outperforms TSSF-nobeta in capturing true cCRE-gene interactions, with improvements observed in 9 out of 18 datasets and declines in 3 datasets. By visualizing the *KLF1* gene (Figure 4d), TSSF-nobeta with KL identifies four active enhancers (TPR=4/6), whereas TSSF-nobeta identifies only three active enhancers (TPR=3/6). Similarly, we used the prior-inspired loss to jointly train TSSF. As shown in Supplementary Figure 4, although TSSF with KL does not show a significant improvement in predictive performance compared to TSSF, its ability to prioritize true cCREs has been effectively enhanced, with improvements observed in 12 out of 20 datasets and declines in 3 datasets. For example, there are three validated enhancers (solid red boxes) surrounding the *BAX* gene. As a result, TSSF fails to identify any of the validated enhancers, but TSSF-KL successfully identifies two of them. Based on the above results, we conclude that the distance-decay prior can further enhance the predictive performance of the two-stage frameworks by aligning attention with regulatory geometry.

### 2.5 Chromatin contacts preferentially rescue distal regulation

Enhancer–gene communication often depends on three-dimensional looping, yet cCRE-centric models rely primarily on linear proximity and local epigenomic profiles. To recover distal regulatory links, we incorporated chromatin contacts into TSSF via two complementary strategies: input augmentation and model augmentation (see **Methods**).

To explore the effectiveness of incorporating chromatin interactions, we applied these two strategies to TSSF, naming it TSSF-LP. Through comparative analysis, we find that although TSSF-LP does not significantly improve the predictive performance compared to TSSF (Figure 5a), it effectively enhances the ability to prioritize true cCREs, with performance improvements observed in 10 out of 19 datasets and performance declines in 4 datasets (Figure 5b). In addition, we performed ablation experiments to investigate the performance of TSSF when using input augmentation or model augmentation, naming them TSSF-LP-in and TSSF-LP-weight, respectively. From the results, TSSF-LP achieves the best predictive performance among the three methods (Figure 5c). TSSF-LP-weight improves the ability to prioritize true cCREs, showing performance gains in 10 out of 18 datasets and declines in 4 datasets (Figure 5d); however, TSSF-LP-in does not achieve significant improvements, showing performance gains in 8 out of 18 datasets and declines in 10 datasets. For example, there are five validated enhancers (solid red boxes) surrounding the *JUNB* gene, including three proximal enhancers and two distal enhancers. As shown in Figure 5e, all models incorporating chromatin interactions successfully identify a distal enhancer, demonstrating the effectiveness of chromatin interactions in capturing distal cCREs

**Figure 5:**
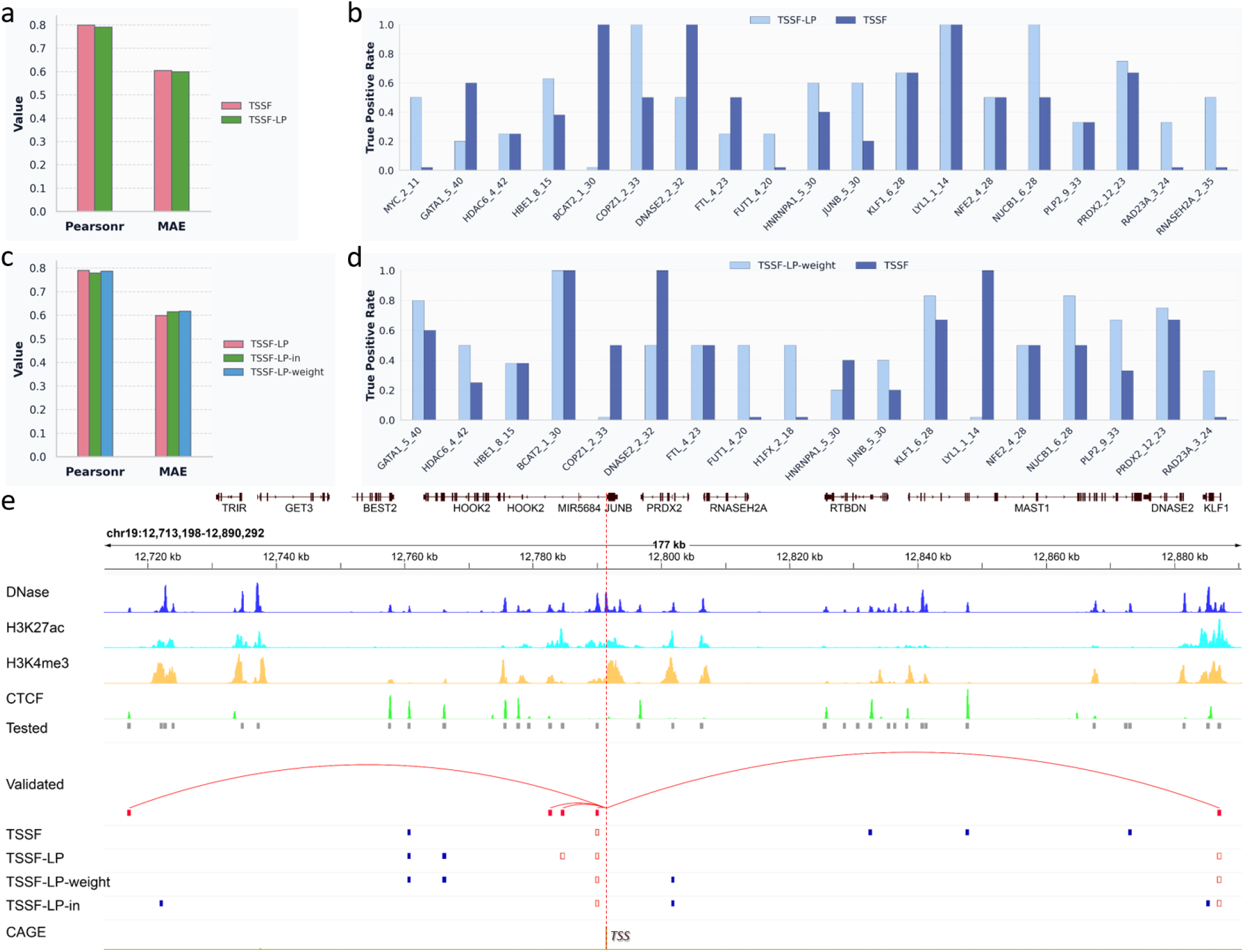
Chromatin interactions facilitate the capture of distal cCRE-gene interactions. (a) Performance comparison (Pearsonr and MAE) between TSSF and TSSF-LP. (b) True positive rates of TSSF and TSSF-LP in identifying genuine cCREs. (c) Performance Comparison of TSSF-LP ablation experiments using input augmentation (TSSF-LP-in) or model augmentation (TSSF-LP-weight). (d) True positive rates of TSSF and TSSF-LP-weight in identifying genuine cCREs. (e) Visualization of *JUNB* gene-active cCREs prioritized by TSSF, TSSF-LP, TSSF-LP-in, and TSSF-LP-weight, where all tested cCREs are represented by solid grey boxes, validated cCREs by solid red boxes, and correctly-identified cCREs by red boxes.

### 2.6 High-attention cCREs recover canonical TF programs

Investigating which transcription factors (TFs) influence a target gene is crucial for gaining a deeper understanding of transcriptional regulation mechanisms. To identify which TFs may regulate the target gene, we applied the augmented-attention strategy to select top-*k* cCREs with high attention scores and divided them into two groups (one group close to genes and another group far from genes). Finally, we utilized the MEME-ChIP and TOMTOM tools to identify highly significant TF motifs.

To analyze the different functions of cCREs surrounding genes, we counted the distribution of types of these top-*k* cCREs. As shown in Figure 6a, the most frequently regulated types of cCREs are proximal elements (e.g., pELS, PLS), and the majority of these elements involve CTCF binding. This observation confirms that promoter-proximal CTCF occupancy sustains distal enhancer-promoter long-range interactions and drives enhancer-dependent transcriptional activation^19^. Furthermore, we identify some key TF motifs through motif analysis (Figure 6b), such as SP1 enriched in proximal regions, AP-1 factors (FOS/JUN family) enriched in distal regions, and CTCF enriched in both proximal and distal regions. For example, SP1 is predominantly a promoter-proximal transcription factor that binds GC-rich elements near transcription start sites to regulate gene expression^20^. AP-1 factors (FOS/JUN family, including inducible JUNB/FOSL1-containing programs) are broadly required for enhancer selection^21^. CTCF is a multifunctional transcription factor that can not only function as a transcriptional activator or repressor by binding to proximal or distal cCREs, but also blocks communication between enhancers and upstream promoters by binding to a transcriptional insulator element^22^.

**Figure 6:**
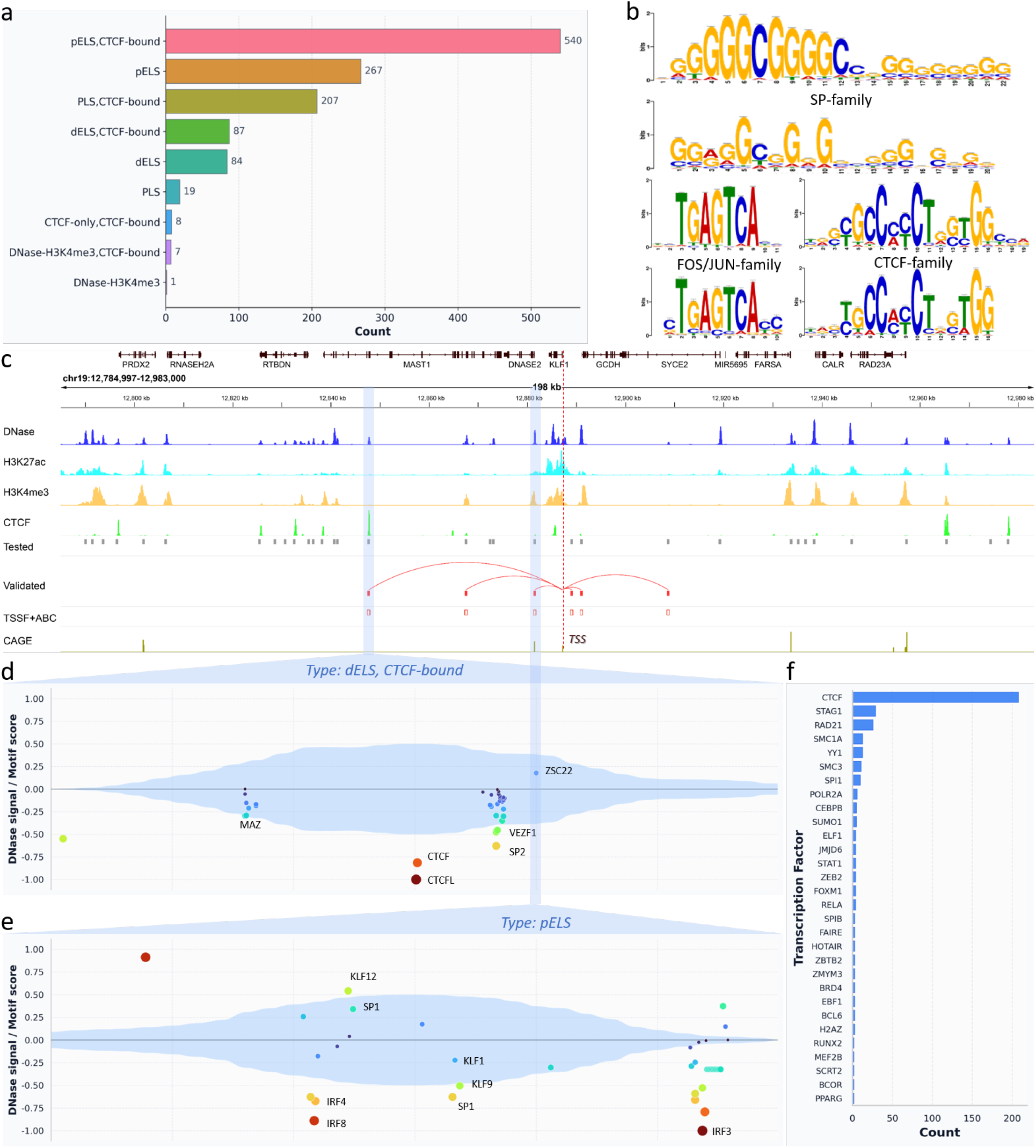
Augmented attention identifies important motifs in specific types of cCREs. (a) Distribution of types of functional cCREs selected by the augmented attention strategy, where *pELS* means proximal enhancer-like signatures, *dELS* means distal enhancer-like signatures, and *PLS* means promoter-like signatures. (b) Motif analysis identifies some important TF motifs in proximal or distal regions. (c) Visualization of *KLF1* gene-active cCREs prioritized by the augmented attention strategy (TSSF+ABC), where all tested cCREs are represented by solid grey boxes, validated cCREs by solid red boxes, and correctly-identified cCREs by red boxes. (d-e) Visualization of the distribution of TF motifs within the two cCREs (type: *dELS, CTCF-bound* and *pELS*), with its core region highlighted by DNase-seq signals; the top and bottom of the panel indicate whether the TF motif is located on the forward or reverse strand, respectively. (f) Distribution of TF ChIP-seq datasets overlapping with these cCREs surrounding the *KLF1* gene.

To further analyze important cCREs surrounding a target gene, we utilized the FIMO tool to identify motif instances with high significance. For example, we picked out two correctly-recognized cCREs from the *KLF1* gene, one with regulatory type dELS, CTCF-bound, and another with regulatory type pELS. By visualizing the distribution of TF motifs on the cCREs (Figures 6c-e), we find that CTCF/CTCFL are the most prominent regulatory factors in the core region of the cCRE (highlighted by DNase-seq signals), exactly matching the function of the cCRE (dELS, CTCF-bound). For another cCRE (labeled pELS), we similarly find that some proximal TFs, such as the SP family, KLF family, and IRF family, are enriched in the core region of the cCRE, which is consistent with the function of the cCRE. Furthermore, we observe that the vast majority of CTCF ChIP-seq peaks are located within these cCREs (Figure 6f), after searching all K562-specific ChIP-seq binding datasets in CistromeDB^23^, confirming that the majority of cCREs involve CTCF binding.

## 3. Discussion

In this study, we developed a two-stage selective framework (TSSF) for gene expression prediction that performs hierarchical refinements: (i) nucleotide selection within each cCRE and (ii) cCRE selection around each target gene. By integrating sequence, DNase-seq, and multimodal epigenomic profiles in a fully Transformer-based architecture, TSSF reduces input redundancy and concentrates model capacity on regulatory features most predictive of transcription. Across 70 human cell types/tissues, TSSF and its lightweight variants (TSSF-signal, TSSF-nobeta) consistently outperform strong baselines in Pearson correlation and MAE, while maintaining favourable computational efficiency. Beyond predictive performance, the two-stage frameworks show improved ability to prioritize functional enhancer–gene interactions in multiple benchmark settings (curated enhancer-gene data, GTEx eQTL-derived pairs, and Hi-ChIP interactions), and the augmented-attention strategy further strengthens recovery of TF motifs in validated enhancers.

This study also provides biologically meaningful extensions. First, a prior-inspired regularization term, motivated by distance-dependent regulatory decay, improves model capability in many datasets and enhances cCRE prioritization even when gains in global expression metrics are modest. Second, incorporating chromatin interaction information improves capture of distal regulatory elements, supporting the value of 3D genome context. Third, motif analyses on high-confidence cCREs recover known regulatory programs (e.g., SP-family motifs in proximal regions, AP-1 in distal regions, and CTCF in both), which increases confidence that the model is learning plausible regulatory mechanisms rather than only statistical correlations. Overall, these results indicate that explicit hierarchical refinements are a promising strategy for improving both predictive performance and interpretability in cCRE-to-gene modeling.

Despite these strengths, several limitations should be acknowledged. First, cross-species generalization remains uneven: performance is better in human-to-mouse transfer than mouse-to-human transfer, indicating that species-specific regulatory grammar and data heterogeneity are not yet fully captured. Second, distal cCRE prioritization remains more challenging than proximal prediction, and improvements are dataset-dependent, suggesting that long-range regulation remains only partially resolved without broader and higher-quality chromatin contact maps. Third, model capability is sensitive to signal quality in key assays (notably DNase and H3K27ac), and noisy or weak signals can lead to underestimation of highly expressed genes. Fourth, several components rely on design assumptions and hand-set hyperparameters (e.g., Beta priors, distance-decay forms, threshold constants), which may affect robustness across data regimes and may require careful retuning for new tissues or species. Fifth, evaluation is largely based on retrospective datasets; additional validation (e.g., CRISPR perturbation benchmarking in unseen contexts) would strengthen causal interpretability. Future work could therefore focus on improving cross-species adaptability, integrating richer 3D and single-cell modalities, and developing more adaptive prior formulations to further enhance robustness and biological interpretability.

## 4. Methods

### Datasets

#### Candidate *cis*-regulatory elements (cCREs)

cCREs for Human were downloaded from the SCREEN Registry Version 3. The Registry contains 1,063,878 human cCREs in GRCh38, which are derived from ENCODE using four assay types (DNase, H3K4me3, H3K27ac, and CTCF). cCREs with DNase-only and low-DNase marks were removed. Given that CREs typically cluster together to form *cis*-regulatory modules (CRMs)^24^, adjacent cCREs within a 600-bp window were merged, reducing the number of cCREs to 726,796.

#### Gene expression

Total RNA-seq and polyA plus RNA-seq data for 70 human cell types or tissues were downloaded from ENCODE. Released gene quantifications mapped to GRCh38 and annotated to GENCODE V29 were retained. Gene expression levels (GEL) were calculated as the average TPM value for each gene, and scaled by the log(1 + *x*) function.

#### Epigenomic tracks

For each cell type or tissue, DNase-seq and ChIP-seq files mapped to GRCh38, including DNase, H3K4me3, H3K27ac, and CTCF types, were downloaded from ENCODE. Multiple read-depth normalized signal files for DNase-seq and fold change over control files for ChIP-seq were retained. Then, multiple signal files for the same type were merged using the *bigWigMerge* software, and scaled by the log_10_(1 + *x*) function.

### Data Construction

In our design concept, we extracted a fixed number of cCREs surrounding genes to predict gene expression levels. Since cCREs are discretely distributed on both sides of genes, we need to integrate DNA sequences and multimodal epigenomic profiles on these cCREs. As shown in Figure 1a, the details are as follows:

#### DNA sequences

DNA sequences of a gene and its surrounding cCREs are expanded to 600bp and transformed into index vectors of shape *l* =600bp following the protocol {‘A’: 0, ‘C’: 1, ‘G’: 2, ‘T’: 3, ‘N’: 4}. Subsequently, these index vectors are individually reshaped to 1 × *l* and assembled along the first dimension to form a two-dimensional matrix of shape (*n* + 1) × *l*, where *n* represents the number of cCREs. For the reverse complement of DNA sequences, we repeat the same process to obtain another two-dimensional matrix of shape (*n* + 1) × *l*.

#### Epigenomic signals

Based on the coordinates of cCREs on the genome, we utilize the *pyBigWig* software to extract epigenomic signals from corresponding epigenomic tracks, including DNase-seq, Histone ChIP-seq (H3K27ac, H3K4me3), and TF ChIP-seq (CTCF). For DNase-seq signals, we employ the same strategy to reshape and organize them into a two-dimensional matrix of shape (*n* + 1) × *l*. For multimodal epigenomic signals, we average nucleotide-resolution signals along each cCRE and organize them into a two-dimensional matrix of shape (*n* + 1) × *d* where *d* = 4 denotes the number of epigenomic tracks.

#### Chromatin loops

Cell-type-specific chromatin loops at 5kb resolution, such as K562 and GM12878, were refined from raw HiChIP data using the FitHiChIP software^25^. If there exist interactions between a gene and its surrounding cCREs, the normalized counts are extracted from chromatin loops and then scaled by the log(1 + *x*) function, otherwise 0. As a result, for each gene, an interaction matrix of shape (*n* + 1) × (*n* + 1) is constructed. Additionally, Hi-C data are processed to calculate contact frequencies between a gene and its surrounding cCREs, following the ABC pipeline method^7^. Then, Hi-C data are scaled using the log(1 + *x*) function and integrated with those multimodal epigenomic profiles.

### Model Design

To reduce computational requirements, we need to extract all cCREs surrounding genes rather than using long-range DNA sequences. For this purpose, we design a two-stage selective framework (TSSF) to improve gene expression prediction (Figure 1b), in which the first stage is to selectively identify key nucleotides by relying on DNA sequences and DNase-seq signals; the second stage is to selectively identify key cCREs by relying on multimodal epigenomic profiles. The selective strategy is based on the information bottleneck theory^17,18^, with the goal of maximizing the mutual information between the compressed representations *Z* and the target variable *Y*, while controlling the information extracted from the input *X*.

Similar to the design concept of Seq2Exp^16^, our framework provides a learnable approach to selectively extract key regulatory elements by leveraging both DNA sequences and epigenomic signals through the information bottleneck mechanism. Based on the objective of information bottleneck (see **Supplementary Note 1**), we define the latent representations as *Z* = *M* ⊙ *X*, where *M* is a soft mask for indicating the importance of each element (such as nucleotide or cCRE), and assume that each selection is independent given an input *X*, i.e., *p*(*M*|*X*) = ∏_*i*_ *p*(*m*_*i*_|*x*_*i*_). After substitution, the objective becomes:

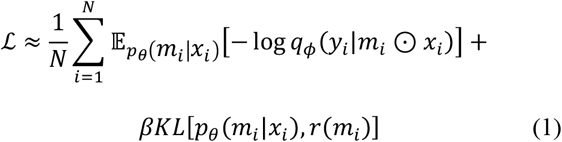

where the first term is a task-specific loss, and the second term imposes a constraint on the learned mask *M*, aligning it with a prior distribution *r*(*M*) without conditioning on any specific input *X*. Based on Equation 1, our core task is to estimate *p*_*θ*_(*M*|*X*), i.e., to learn the mask by incorporating input information.

#### Two-stage Selective Framework

Our framework comprises two stages for selecting key regulatory elements: the first stage uses DNA sequences and DNase-seq signals to identify key nucleotides, and the second stage employs multimodal epigenomic profiles to identify key cCREs. Accordingly, we need to estimate two independent masks based on DNA sequences and epigenomic signals, i.e., 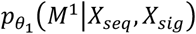 and 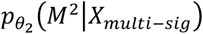

Given that an input *X* consists of DNA sequences *X*_*seq*_ and DNase-seq signals *X*_*sig*_, we first need to estimate 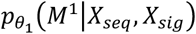 to generate a mask *M*^1^ for identifying key nucleotides. We assume that, conditioned on the selection of regulatory nucleotides *M*^1^, DNA sequences and DNase-seq signals are conditionally independent of each other, i.e.,

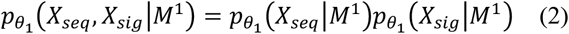

By applying Bayes’ theorem, the estimation of 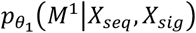 can be further decomposed into two terms involving *X*_*seq*_ and *X*_*sig*_. Specifically, we have:

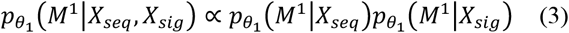

where the two terms represent the contributions from DNA sequence and DNase-seq signals, respectively. As a result, we can independently estimate 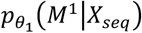 and 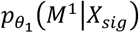, which simplifies the estimation of 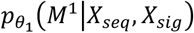.

To specify the distribution of *p*_*θ*_(*M*|*X*), we choose the Beta distribution based on the following desirable properties: (i) Beta distribution can typically quantify success rates^26^, with parameters α and β representing the weights for selecting and not selecting nucleotides, respectively, i.e., when α > β, these nucleotides are more likely to be selected, and vice versa; (ii) the product of two Beta distributions remains a Beta distribution, which simplifies subsequent mathematical calculations and provides a consistent fitting objective for models. Therefore, we assume that 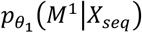 follows a Beta distribution with parameters α_*seq*_ and β_*seq*_, and that 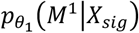 follows a Beta distribution with parameters α_*sig*_ and β_*sig*_. After the product of these two Beta distributions, we obtain a joint Beta distribution with parameters:

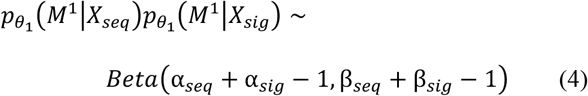

where α_*seq*_ and β_*seq*_ could be estimated adaptively by a deep learning model, while α_*sig*_ and β_*sig*_ could be determined by DNase-seq signals.

From the objective in Equation 1, the KL divergence between 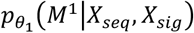 and *r*(*M*^1^) needs to be computed. To simplify the calculation process, we assume that the prior distribution *r*(*M*^1^) follows a Beta distribution as well. Therefore, we have 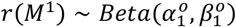, where 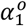 and 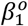 are related to the sparsity of the mask *M*^1^ and are empirically set in experiments.

Importantly, we can estimate the mask *M*^1^ only from DNase-seq signals, i.e., 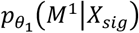, thereby reducing model complexity and computational requirements. Since these signals directly reflect the regulatory states of nucleotides, higher values indicate greater openness and thus a higher likelihood of being involved in regulatory activities. Therefore, the estimation of the mask *M*^1^ could be simplified from 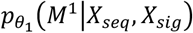 to 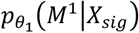.

Secondly, given multimodal epigenomic profiles, including DNase-seq, Histone ChIP-seq (H3K27ac, H3K4me3), and CTCF ChIP-seq, we need to estimate 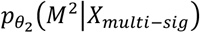 to generate a mask *M*^2^ for identifying key cCREs. Here *X*_*multi*−*sig*_ denotes the per-cCRE profile vector obtained by averaging nucleotide-resolution epigenomic signals within each cCRE, representing the global cCRE context, i.e., *X*_*multi*−*sig*_ = {*X*_*DNase*_, *X*_*H*3*K*27*ac*_, *X* _*H*3*K*4*me*3_, *X* _*CTCF*_}. Similar to the first stage, we assume that 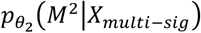 also follows a Beta distribution with parameters α_*multi*−*sig*_ and β_*multi*−*sig*_.

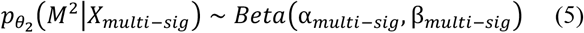

where *d* denotes the number of epigenomic tracks, and 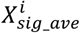 denotes the average value of the *i*-th epigenomic data. The parameters α_*multi*−*sig*_ and β_*multi*−*sig*_ could be determined by multimodal epigenomic profiles.

Similarly, to compute the KL divergence between 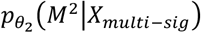 and *r*(*M*^2^), we assume that the prior distribution *r*(*M*^2^) follows a Beta distribution as well. Therefore, we have 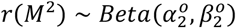, where 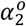 and 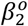 are related to the sparsity of the mask *M*^2^ and are empirically set in experiments. Note that 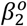 increases exponentially with distance, i.e., (1 − *exp*(−*r* * *D*)) * *D*, where *r* denotes the decay rate and *D* denotes the distance between cCREs and genes, indicating a higher threshold for selecting distal cCREs. This assumption is consistent with a prior found in this study that the influence of cCREs on genes decreases with distance.

Finally, the objective of TSSF becomes:

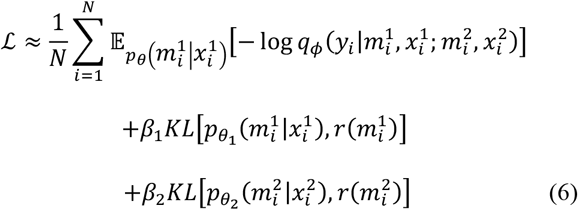

where 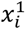 denotes DNA sequences and DNase-seq signals, and 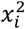 denotes multimodal epigenomic profiles; 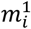 and 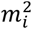 denote two masks for selecting key nucleotides and cCREs in the *i*-th sample, respectively. The first term is a task-specific loss, such as Smooth L1 Loss, and *β*_1_, *β*_2_ are hyperparameters that specify the contribution of the *KL* loss to the prediction.

#### Model Architecture

As shown in Figure 1b, our proposed model comprises two stages for selectively extracting relevant nucleotides and cCREs. The first stage generates a mask distribution 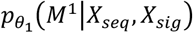 from DNA sequences and DNase-seq signals, through a Beta Transformer, which is then used to extract critical nucleotide-level information, i.e., (*X*_*seq*_ + *X*_*sig*_) ⨀ *M*^1^. Subsequently, a Site Transformer is employed to learn hidden representations *X*_*hidden*_ from the filtered input. The second stage generates a mask distribution 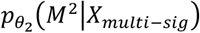 from multimodal epigenomic profiles, which is then used to extract critical cCRE-level information, i.e., (*X*_*multi*−*sig*_ + *X*_*hidden*_) ⨀ *M*^2^. Subsequently, a cCRE Transformer is employed to predict gene expression levels (GEL) by integrating all features learned from cCREs. The details are as follows:

The Beta Transformer, consisting of a two-layer T5 Transformer and a linear projection layer, utilizes DNA sequences to predict the parameters α_*seq*_ and β_*seq*_ for 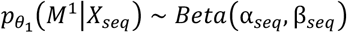. Specifically, we have:

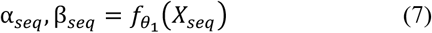

where the Beta Transformer outputs the *l* -dimensional parameters α_*seq*_ and β_*seq*_, with *l* being the length of DNA sequences. Each position in the sequence is associated with a pair of values, which parameterize the Beta distribution for that nucleotide.

For the parameters α_*sig*_ and β_*sig*_, we follow the intuition that higher signals increase the likelihood of selecting corresponding nucleotides. Therefore, we directly use the epigenomic signals as the parameter α_*sig*_, which controls the selection weight for each nucleotide. The parameter β_*sig*_, which represents a selection threshold, is set to a constant. Specifically, we have:

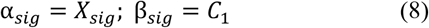

With these settings, we make the learning of α_*sig*_ and β_*sig*_ non-parametric while maintaining the influence of signal strength without introducing additional learnable parameters. Note that we can estimate the mask distribution 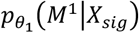 only from DNase-seq signals, thereby reducing model complexity and computational requirements.

After estimating these parameters, the mask *M*^1^ is sampled from a joint Beta distribution according to Equation 4. Its values range from 0 to 1, representing the probability of each nucleotide being selected. If only DNase-seq signals are used, the mask *M*^1^ is sampled from a Beta distribution with the parameters α_*sig*_ and β_*sig*_.

The Site Transformer, consisting of a two-layer T5 Transformer, utilizes the filtered input (*X*_*seq*_ + *X*_*sig*_) ⨀ *M*^1^ to produce a hidden representations *X*_*hidden*_.

Similarly, following the intuition that higher epigenomic signals increase the likelihood of selecting corresponding cCREs, we integrate multimodal epigenomic profiles to estimate the parameters α_*multi*−*sig*_ and β_*multi*−*sig*_. Therefore, we directly use the geometric mean of all epigenomic signals as the parameter α_*multi*−*sig*_, which controls the selection weight for each cCRE. The parameter β_*multi*−*sig*_, representing a selection threshold, is set as a constant. Specifically, we have:

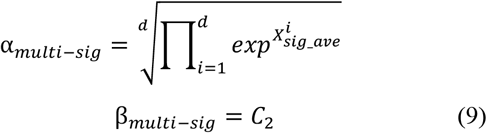

With these settings, we make the learning of α_*multi*−*sig*_ and β_*multi*−*sig*_ non-parametric. Then, the mask *M*^2^ is sampled from a Beta distribution with the parameters α_*multi*−*sig*_ and β_*multi*−*sig*_. Its values range from 0 to 1, representing the probability of each cCRE being selected.

The cCRE Transformer, consisting of a four-layer T5 Transformer and a prediction module, utilizes the filtered hidden representations (*X*_*multi*−*sig*_ + *X*_*hidden*_)⨀*M*^2^ to predict gene expression levels (GEL). Specifically, the T5 Transformer first generates a matrix *Z* of shape (*n* + 1) × *h*. Then, the gene row representing the associations between a gene and its surrounding cCREs is extracted from this matrix *Z*, and fused with the mRNA half-life features^27^. Finally, the fused features are fed into the prediction module, a fully connected feed-forward network with a Softplus activation, to predict gene expression levels.

#### T5 Transformer

The Beta Transformer, Site Transformer, and cCRE Transformer all adopt the T5 Transformer architecture^28^, in which the Beta Transformer is used to estimate the parameters of Beta distribution, the Site Transformer is used to learn important information from cCREs, and the cCRE Transformer is used to capture the associations between genes and their surrounding cCREs.

Typically, each block in the T5 Transformer consists of a multi-head self-attention layer and a position-wise feed-forward network. In each self-attention layer, scaled dot-product attentions are performed as follows: the query *Q*, key *K*, and value *V* are calculated through a linear projection of the input *X*, respectively; the attention weights are calculated by 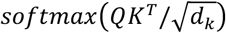 where *d*_*k*_ denotes the number of channels; the value *V* representing the semantics of all embeddings is aggregated according to the attention weights.

For position embeddings, we apply a relative positional embedding 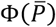 onto the attention weights, where 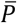 represents the relative position between nucleotides within DNA sequences in the Beta Transformer and Site Transformer, while in the cCRE Transformer, 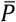 represents the relative distance between genes and their surrounding cCREs.

Moreover, we mask the attention weights assigned to genes themselves to encourage the model to focus on capturing the associations between genes and their surrounding cCREs. Finally, the feed-forward network integrates the outputs from multi-head attention layers and introduces non-linearity. The calculation process can be described as follows:

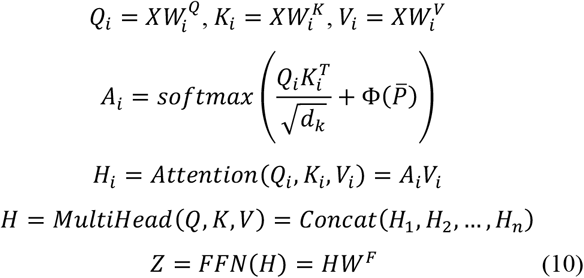

where 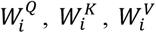 represent the weights of three linear layers, respectively, and *H*_*i*_ denotes the outputs from the *i*-th self-attention layer, and *W*^*F*^ denotes the weights of the feed-forward network.

#### Model Optimization

After obtaining the core parameters α and β of all Beta distributions, we generate the masks *M*^1^ and *M*^2^ by sampling from their corresponding distributions. To train the loss function defined in Equation 6, we treat the Beta distribution as a special case of the Dirichlet distribution, and achieve differentiable sampling from the Dirichlet distribution through the re-parameterization technique^29^. This technique guarantees that the sampling process remains differentiable with respect to the parameters α and β, enabling efficient backpropagation and optimization.

During inference, we directly use the expected value of the Beta distribution as the mask rather than sampling from it. For a Beta distribution with parameters α and β, its expected value 𝔼[*M*] = α/(*α* + *β*) provides a deterministic and efficient way to generate the mask without introducing randomness.

#### Implementation Details

During data splitting, the coding genes from Chromosome 16 were used to validate the models, the coding genes from Chromosomes 8 and 9 to test the models, and the coding genes from the remaining Chromosomes, except Chromosome Y, to train the models.

During training, we employed a ‘warm-up’ strategy to mitigate the impact of parameter initialization. After multiple warm-up rounds, we selected the best-performing model to continue training. All models were trained using the AdamW algorithm, with the initial learning rate gradually increased from 1e-06 to 1e-03. Then, the models were validated every 1000 steps during 15000 training steps. All models were implemented using PyTorch and trained on a single A100 GPU.

### Prior-inspired Loss

In this study, we found a prior that the regulatory effect of cCREs on target genes diminishes with increasing distance, similar to a normal-like distribution. Inspired by this prior, we designed a loss function based on the KL divergence between the prior and attention weights, where the prior is defined as an exponential decay function. By minimizing the KL divergence, we can guide the distribution of attention weights to converge toward the distribution of the decay function during training. Finally, we added the prior-inspired loss term to the model loss to jointly train our proposed models.

Specifically, (i) the multi-head attention weights were extracted from transformer blocks and then averaged across all heads and layers, denoted by *Ā*(ii) the gene rows of *Ā* were extracted, denoted by *Ā*_*g*_; (iii) the decaying function *D*_*g*_ was simulated with an exponential function; (iv) after that, *D*_*g*_ and *Ā*_*g*_ were normalized to a range of 0 to 1; (v) only those with gene expression levels greater than zero were selected for calculating the KL divergence, e.g., *D*_*g*>0_ and *Ā*_*g* > 0_.

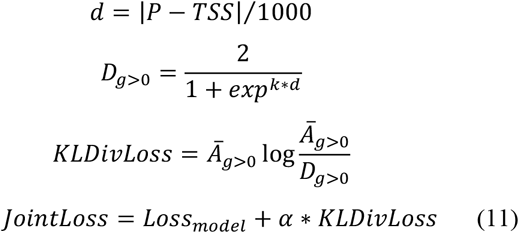

where *P* denotes the position of cCREs in the genome, *TSS* represents the transcription start site, and *d* indicates the distance between a gene and its surrounding cCREs. *α* is a hyperparameter used to specify the relative contribution of the two losses.

### Cell-type Specific Gene Prediction

To evaluate the model’s performance in predicting cell type-specific gene expression, we followed the steps below to select cell type-specific genes from the K562 and GM12878 cell lines: the log_2_FC value for each gene was calculated using the formula: log_2_(1 + *TPM*^*K*562^) − log_2_(1 + *TPM*^*GM*12878^) ; (ii) the genes with log_2_ *FC* > 1 were considered as K562-specific, while those with log_2_ *FC* < −1 were considered as GM12878-specific. After training the two cell-type-specific models, we performed corresponding cross-cell-type prediction.

### Incorporation of Chromatin Interactions

Genome-wide chromatin interactions, such as Hi-C and HiChIP, may provide valuable information for guiding models to capture the long-range regulatory relationships between genes and cCREs. To incorporate chromatin interactions, we therefore applied two strategies: input augmentation and model augmentation.

#### Input augmentation

We integrated the contact frequencies between cCREs and genes with multimodal epigenomic profiles, directly serving as input for the second stage of our proposed framework, i.e., *X*_*multi*−*sig*_ = {*X*_*DNase*_, *X*_*H*3*K*27*ac*_, *X*_*H*3*K*4*me*3_, *X*_*CTCF*_, *X*_*contact*_}.

#### Model augmentation

We first refined several significant chromatin interactions from raw HiChIP data using the FitHiChIP software^25^ and transformed them into an interaction matrix *M*. After that, we put the interaction matrix *M* directly into the self-attention layers of the cCRE Transformer with the purpose of increasing the attention weights of physical cCRE-gene interactions. Through this method, the two-stage frameworks are guided to pay more attention to physical cCRE-gene interactions. Correspondingly, the calculation of *A*_*i*_ in Equation 10 is modified as follows:

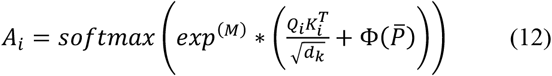

### cCRE Prioritization

Linking candidate enhancers to their target genes through high-throughput experiments remains a significant challenge in terms of both time and cost. Against this backdrop, accurately prioritizing candidate enhancer-gene interactions through computational methods has become increasingly important. To achieve this goal, we applied three methods: attention weights and augmented attention weights.

#### Attention weights

After training the models, we extracted the multi-head attention weights from the cCRE Transformer and subsequently performed element-wise averaging across all heads and layers. To focus on the attention between the gene and its surrounding cCREs, we extracted the gene row from the attention matrix and normalized it to a range of 0 to 1. Thus, by matching the query index at cCRE locations, we can directly observe the importance of each cCRE to that gene.

#### Augmented attention weights

After obtaining the attention weights, we can calculate the augmented attention weights by multiplying the ABC scores^7^ by the attention weights.

### Motif Analysis

To identify key TF motifs that may regulate gene expression, we performed motif analysis using high-confidence cCREs estimated from the augmented attention weights. Specifically, (i) we used the two-stage framework TSSF to predict gene expression levels on the test set and then selected true positives (TPs) from these predictions (see **Supplementary Note 2**); (ii) we applied the augmented attention weights to calculate attention scores between all cCREs and genes, from which we selected top-*k* cCREs (*k*=5); (iii) we divided these cCREs into two groups based on distance: one group close to genes (<10 kb) and another group far from genes (>10 kb); (iv) we performed comprehensive motif analysis on both groups using the MEME-ChIP tool^30^; (v) we utilized the TOMTOM tool^31^ to identify highly significant TF motifs by matching against public motif databases.

To conduct an in-depth analysis of some specified cCREs, we first selected those cCREs that were correctly prioritized by the augmented attention weights, and then utilized the FIMO tool^32^ to identify motif instances with high significance (p-value < 1e-06). To highlight the TF motif distribution in the core regions (i.e., regions with high DNase signals), we visualized these motif instances within these cCREs alongside their corresponding DNase-seq signals.

### Model Comparisons

To benchmark our proposed methods against existing methods, we conducted comparative experiments involving four state-of-the-art models (CREaTor^13^, EPInformer^14^, ScPGE^15^, and Seq2Exp^16^) and three TSSF variants (TSSF, TSSF-signal, and TSSF-nobeta).

Specifically, CREaTor is a deep neural network based on hierarchical attention that utilizes cCREs within open chromatin regions, combined with ChIP-seq data of TFs and histone modifications, to predict gene expression levels. EPInformer is a Transformer-based framework that predicts gene expression by integrating DNA sequences, epigenomic signals, and promoter-enhancer interactions. ScPGE is a scalable computational framework that predicts gene expression by assembling DNA sequences, TF binding scores, and epigenomic tracks of discrete cCREs into 3-dimensional tensors. Seq2Exp is a sequence-to-expression network designed to explicitly discover and extract regulatory elements that drive target gene expression, thereby enhancing the accuracy of the gene expression prediction.

TSSF combines DNA sequence, DNase-seq, and multimodal epigenomic profiles in a two-stage Transformer framework, using Beta-distributed masks to refine nucleotide- and cCRE-level representations. TSSF-signal retains the same two-stage design but estimates nucleotide-level masks in stage 1 from DNase-seq alone. TSSF-nobeta removes explicit Beta masking at both stages and instead relies on standard attention to adaptively weight nucleotides and cCREs.

Following the official guide manual, we employed default hyperparameter settings to train CREaTor, EPInformer, ScPGE, and Seq2Exp from scratch, covering 70 human cell lines or tissue samples. Due to the unavailability of chromatin loops in most datasets, we excluded chromatin loops during model training.

### cCRE-gene Interactions

We collected different types of cCRE-gene interactions to evaluate the model’s performance in prioritizing important cCREs for target genes.

#### Enhancer-gene dataset

Three K562-specific enhancer-gene interaction datasets were collected from Fulco *et al*., Gasperini *et al*., and Schraivogel *et al*., where enhancer-gene pairs were categorized into four distance-based groups: ‘<10kb’, ‘10-50kb’, ‘50-100kb’, and ‘>100kb’. For all three datasets, the genomic coordinates of candidate enhancers were first converted from hg19 to hg38 using the LiftOver software, then integrated by gene names.

#### GTEx dataset

Three eQTL fine-mapping datasets were collected from GTEx v10, involving ‘EBV-transformed Lymphocytes’, ‘Whole Blood’, and ‘Liver’. From each eQTL fine-mapping dataset, we identified causal SNPs as positive SNPs with a posterior inclusion probability (PIP) > 0.9 and an absolute slope > 0.5. On the contrary, we identified non-causal SNPs as negative SNPs with a PIP < 0.001 and an absolute slope > 0.5. Similarly, all interactions were categorized into four distance-based groups.

#### Hi-ChIP dataset

K562-specific Hi-ChIP data (1-kb resolution) targeting H3K27ac were collected from HiChIPdb, representing candidate cCRE-gene interactions. The data processing workflow is as follows: (i) the genomic coordinates of read pairs were converted from hg19 to hg38 using the LiftOver software; (ii) all significant interactions (*q*-value < 1e-15 and count > 30) within 100 kb were regarded as positive samples according to the rule: read pairs intersect with both cCREs and active genes, where active genes are defined as those with expression levels above the 3/4 percentile of all expression levels; (iii) candidate cCREs within 100kb of target genes were randomly selected from all cCREs as negative samples. Similarly, all interactions were categorized into four distance-based groups.

## Software Availability

The two-stage frameworks were implemented in PyTorch. The source code is available at https://github.com/turningpoint1988/TSSF.

## Acknowledgments

This work was supported in part by the National Natural Science Foundation of China (Nos. 62372255), and partly supported by the Natural Science Foundation of Zhejiang Province (No. LMS25F020001).

## Author Contributions

D.S. Huang and Q.H. Zhang conceived the basic idea; Q.H. Zhang and M. Xing designed specific experiments, developed methods, and wrote the manuscript; Z.P. Li carried out the data analysis and model interpretation; Q. Liao provided some advice for writing the manuscript.

## Conflict of Interest

The authors declare no conflict of interest.

## Notes

### Competing Interest Statement

The authors have declared no competing interest.

## References

1 Furlong, E. E. & Levine, M. Developmental enhancers and chromosome topology. Science 361, 1341–1345 (2018).

2 Nasser, J. et al. Genome-wide enhancer maps link risk variants to disease genes. Nature 593, 238–243 (2021).

3 Consortium, G. The GTEx Consortium atlas of genetic regulatory effects across human tissues. Science 369, 1318–1330 (2020).

4 Yazar, S. et al. Single-cell eQTL mapping identifies cell type–specific genetic control of autoimmune disease. Science 376, eabf3041 (2022).

5 Rao, S. S. et al. A 3D map of the human genome at kilobase resolution reveals principles of chromatin looping. Cell 159, 1665–1680 (2014).

6 Mumbach, M. R. et al. HiChIP: efficient and sensitive analysis of protein-directed genome architecture. Nature methods 13, 919–922 (2016).

7 Fulco, C. P. et al. Activity-by-contact model of enhancer– promoter regulation from thousands of CRISPR perturbations. Nature genetics 51, 1664–1669 (2019).

8 Zhou, J. & Troyanskaya, O. G. Predicting effects of noncoding variants with deep learning–based sequence model. Nature methods 12, 931–934 (2015).

9 Zhou, J. et al. Deep learning sequence-based ab initio prediction of variant effects on expression and disease risk. Nature genetics 50, 1171–1179 (2018).

10 Kelley, D. R. Cross-species regulatory sequence activity prediction. PLoS computational biology 16, e1008050 (2020).

11 Avsec, Ž. et al. Effective gene expression prediction from sequence by integrating long-range interactions. Nature methods 18, 1196–1203 (2021).

12 Karbalayghareh, A., Sahin, M. & Leslie, C. S. Chromatin interaction–aware gene regulatory modeling with graph attention networks. Genome Research 32, 930–944 (2022).

13 Li, Y. et al. CREaTor: zero-shot cis-regulatory pattern modeling with attention mechanisms. Genome Biology 24, 266 (2023).

14 Lin, J., Li, Z., Zhao, Y., Luo, R. & Pinello, L. EPInformer: scalable and integrative prediction of gene expression from promoter-enhancer sequences with multimodal epigenomic profiles. Nature Communications 17, 3975 (2026).

15 Zhang, Q. et al. A scalable computational framework for predicting gene expression from candidate cis-regulatory elements. Genome Research (2026).

16 Su, X., Yu, H., Zhi, D. & Ji, S. In 13th International Conference on Learning Representations, ICLR 2025. 58104-58119 (International Conference on Learning Representations, ICLR).

17 Alemi, A. A., Fischer, I., Dillon, J. V. & Murphy, K. Deep variational information bottleneck. arXiv preprint 1612.00410 (2016).

18 Belinkov, Y. & Henderson, J. In International Conference on Learning Representations.

19 Kubo, N. et al. Promoter-proximal CTCF binding promotes distal enhancer-dependent gene activation. Nature structural & molecular biology 28, 152–161 (2021).

20 Hasegawa, Y. & Struhl, K. Different SP1 binding dynamics at individual genomic loci in human cells. Proceedings of the National Academy of Sciences 118, e2113579118 (2021).

21 Vierbuchen, T. et al. AP-1 transcription factors and the BAF complex mediate signal-dependent enhancer selection. Molecular cell 68, 1067-1082. e1012 (2017).

22 Kim, S.Yu, N.-K. & Kaang, B.-K. CTCF as a multifunctional protein in genome regulation and gene expression. Experimental & Molecular Medicine 47, e166–e166, doi:10.1038/emm.2015.33 (2015).

23 Zheng, R. et al. Cistrome Data Browser: expanded datasets and new tools for gene regulatory analysis. Nucleic acids research 47, D729–D735 (2019).

24 Hardison, R. C. & Taylor, J. Genomic approaches towards finding cis-regulatory modules in animals. Nature Reviews Genetics 13, 469–483 (2012).

25 Bhattacharyya, S., Chandra, V., Vijayanand, P. & Ay, F. Identification of significant chromatin contacts from HiChIP data by FitHiChIP. Nature Communications 10, 4221, doi:10.1038/s41467-019-11950-y (2019).

26 Edition, S. Bayesian data analysis. (CRC press, 2013).

27 Agarwal, V. & Shendure, J. Predicting mRNA abundance directly from genomic sequence using deep convolutional neural networks. Cell reports 31 (2020).

28 Raffel, C. et al. Exploring the limits of transfer learning with a unified text-to-text transformer. Journal of machine learning research 21, 1–67 (2020).

29 Figurnov, M., Mohamed, S. & Mnih, A. Implicit reparameterization gradients. Advances in neural information processing systems 31 (2018).

30 Machanick, P. & Bailey, T. L. MEME-ChIP: motif analysis of large DNA datasets. Bioinformatics 27, 1696–1697 (2011).

31 Gupta, S., Stamatoyannopoulos, J. A., Bailey, T. L. & Noble, W. S. Quantifying similarity between motifs. Genome biology 8, R24 (2007).

32 Grant, C. E., Bailey, T. L. & Noble, W. S. FIMO: scanning for occurrences of a given motif. Bioinformatics 27, 1017–1018 (2011).

33 Reyna, J. et al. Loop Catalog: a comprehensive HiChIP database of human and mouse samples. Genome Biology 26, 175 (2025).

