## Supplemental Notes and Figures for "Hierarchical refinements of cis-regulatory inputs improve scalable gene expression prediction"

### Supplementary Notes:

#### Note 1: Description of Information Bottleneck

The information bottleneck method is a widely used technique in machine learning on tasks for images (Chen et al., 2018), language data (Belinkov et al., 2020; Lei et al., 2016; Paranjape et al., 2020; Bastings et al., 2019; Jain et al., 2020) or graph data (Wu et al., 2020; Miao et al., 2022). Its goal is to maximize the mutual information between compressed representations $Z$ and the target variable $Y$, expressed as $I(Z;Y)$, while controlling the information extracted from the input $X$. A straightforward approach would be to set $Z = X$, but this retains the full complexity of $X$, which makes the optimization process challenging.

To address this, researchers impose a constraint on the information transferred from $X$ to $Z$, ensuring that $I(X;Z) \leq I_c$, where $I_c$ is an information constraint that allows us to capture only the most critical compressed representations. The information bottleneck objective becomes maximizing:

$$\mathcal{L} = I(Z;Y) - \beta I(X;Z) \quad (1)$$

where β is a hyperparameter that balances the trade-off between compression and relevance. However, directly optimizing this objective is challenging. To overcome this, Chen et al. (2018) proposes to maximize a lower bound approximation, which leads to minimizing the following expression:

$$\mathcal{L} \approx \frac{1}{N}\sum_{i=1}^{N} \mathbb{E}_{p_\theta(Z|x_i)}\left[-\log q_\phi(y_i|Z)\right] + \beta KL[p_\theta(Z|x_i), r(Z)] \quad (2)$$

where $p_\theta(Z|x_i)$ is a parametric approximation of $Z$, $q_\phi(y_i|Z)$ is a variational approximation of the true distribution $p(y_i|Z)$, and $r(Z)$ approximates the marginal distribution $p(Z)$.

#### Note 2: Categorization of predictions

To understand the different patterns of false positives (FPs), true positives (TPs), false negatives (FNs), and true negatives (TNs), the predictions from the test set were categorized into FPs, TPs, FNs, and TNs by following the two steps: (i) the predicted and true gene expression levels were converted to a range of 0 to 1 by the equation: $(x - x_{min})/(x_{max} - x_{min})$; (ii) we defined as FPs by the rule: the prediction (*P*) is above 0.7 and the true (*T*) is below 0.2, as TPs by the rule: both *P* and *T* are above 0.7, as FNs by the rule: *P* is above 0.2 and *T* is below 0.7, and as TNs by the rule: both *P* and *T* are below 0.2.

### Supplementary Figures:

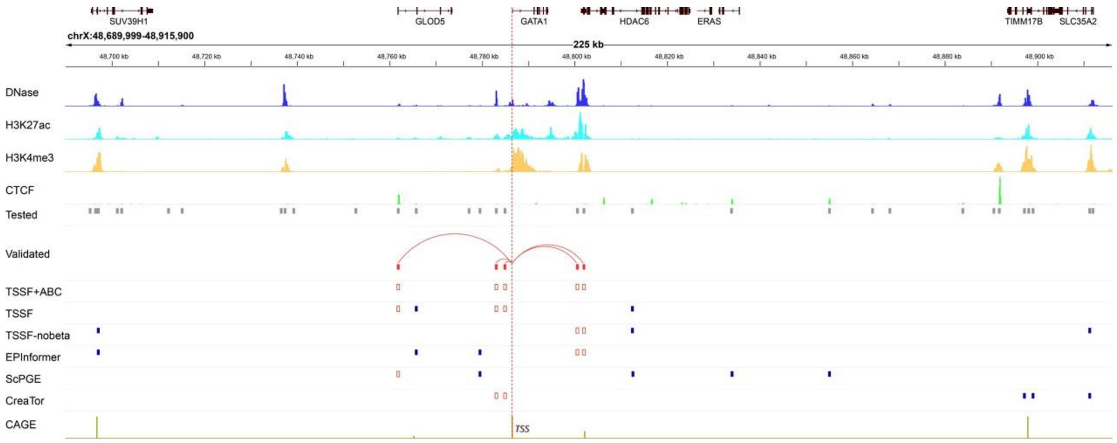


**Figure 1:** Visualization of *GATA1* gene-active cCREs prioritized by multiple methods, where all tested cCREs are represented by solid grey boxes, validated cCREs by solid red boxes, and correctly-identified cCREs by red boxes.

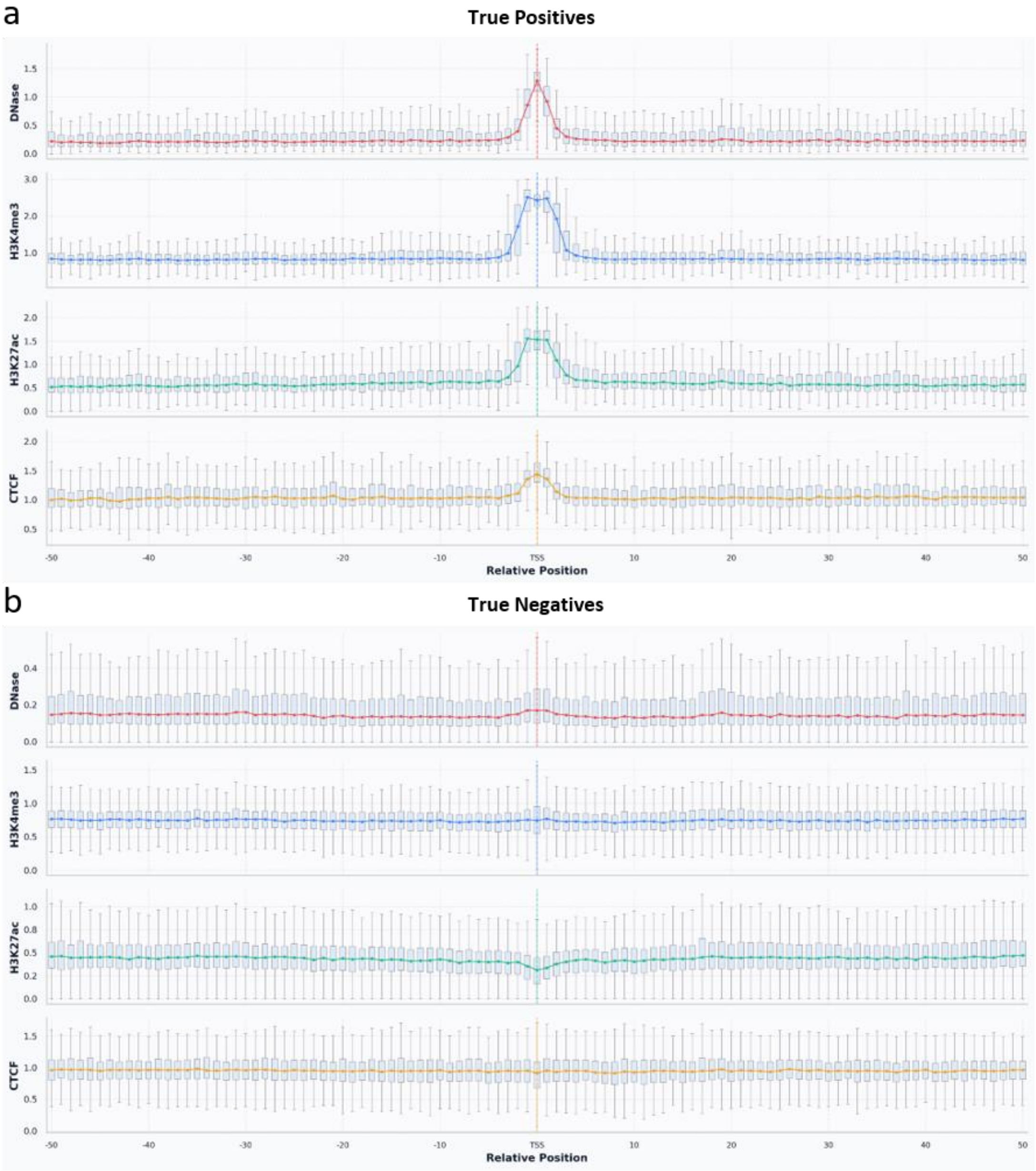


**Figure 2: Different patterns found in true positives (TPs) and true negatives (TNs).** (a) A pattern found in TPs that the regulatory effect of cCREs on target genes diminishes with increasing distance. (b) A pattern found in TNs that the distribution is almost linear, with DNase and H3K27ac signals approaching zero.

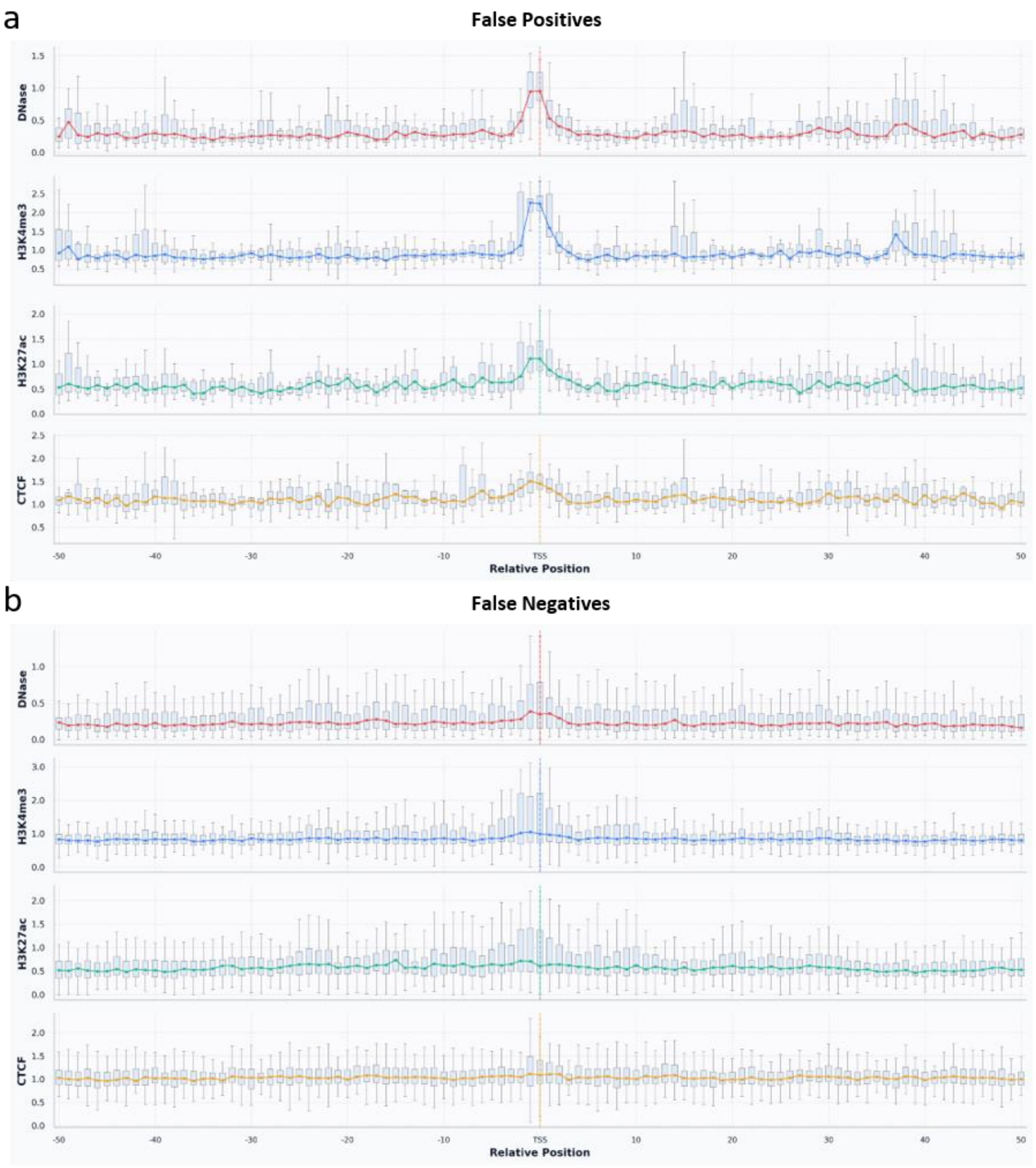


**Figure 3:** (a) A pattern found in FPs that the distribution of four chromatin signals shows a normal-like distribution similar to that of TPs. (b) A pattern found in FNs that the distribution is almost linear similar to that of TNs.

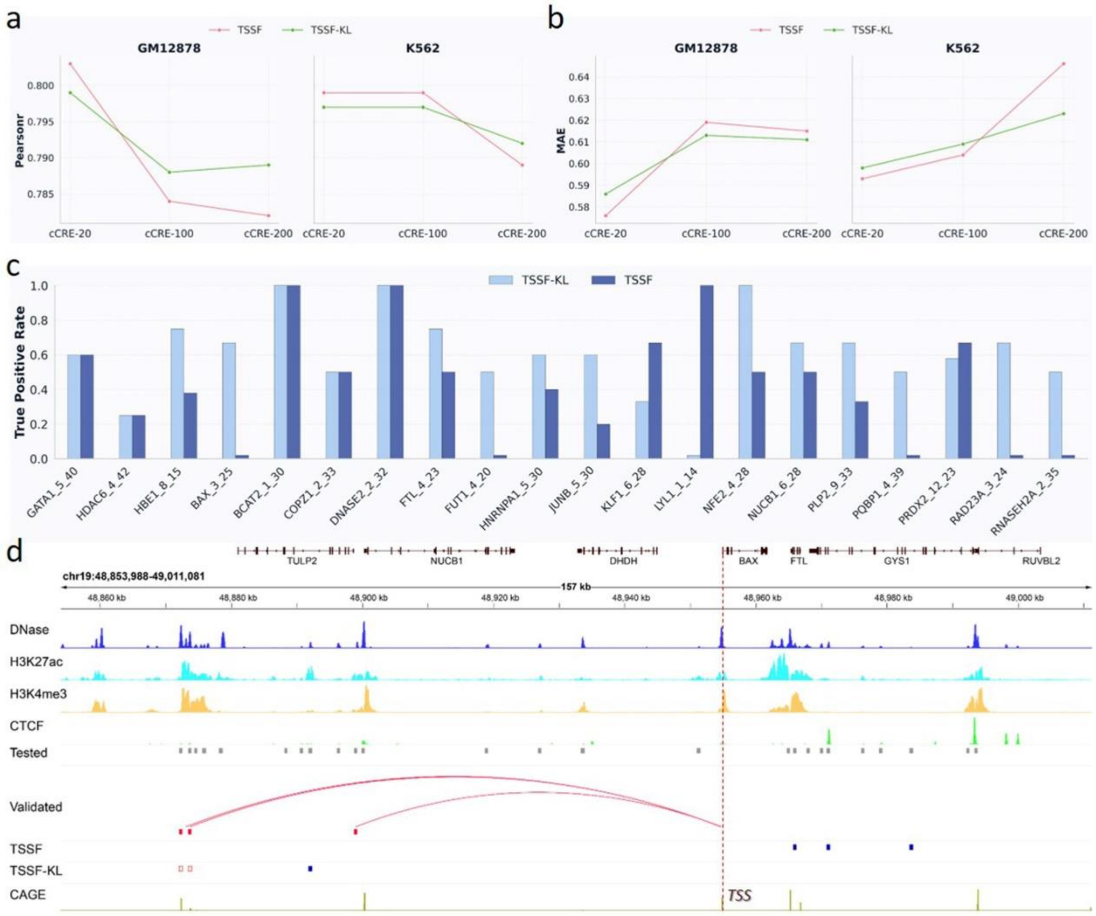


**Figure 4:** (a-b) Performance comparison (Pearsonr and MAE) between TSSF and TSSF-KL. (c) True positive rates of TSSF and TSSF-KL in identifying genuine cCREs. (d) Visualization of *BAX* gene-active cCREs prioritized by TSSF and TSSF-KL, where all tested cCREs are represented by solid grey boxes, validated cCREs by solid red boxes, and correctly-identified cCREs by red boxes.
